# The FXR agonist obeticholic acid does not stimulate liver regeneration in hepatectomized mice

**DOI:** 10.1101/2022.11.10.515905

**Authors:** Kim M.C. van Mierlo, Carola Dahrenmoller, Valérie Lebrun, Peter L.M. Jansen, Cornelis H.C. Dejong, Isabelle A. Leclercq, Steven W.M. Olde Damink, Frank G. Schaap

## Abstract

**Background:** Postresectional liver failure (PLF) is a dreaded complication after partial hepatectomy (PH). Data from animal experiments indicate that endogenous ligands (*i.e*. bile salts) can stimulate liver regeneration and prevent liver injury after PH, via hepatic Fxr and the ileal Fxr-Fgf15 axis.

**Aim:** To investigate whether exogenous activation of the Fxr pathway with the semi-synthetic bile acid derivative obeticholic acid (OCA) could stimulate postresectional liver regeneration in mice.

**Methods:** Twelve weeks old male C57BL6/J mice were pre-treated with OCA (10 mg/kg/day) or vehicle, and after 7 days subjected to 70% PH. Mice were sacrificed at 24, 48 and 72 hrs after PH, and liver injury, secretory function, and regenerative indices were assessed. In a second study, OCA pre-treated mice received oral sucrose supplementation in the postoperative trajectory, and a group of mice receiving intraperitoneal injection of FGF19 was included as a positive control group. Here, mice were sacrificed at 48 hours after PH.

**Results:** No effect could be detected on liver mass recovery after PH, although responses of *Cyp7a1*, *Cyp8b1* and other Fxr target genes implied general effectiveness of OCA treatment. OCA had no consistent effects on the number of Ki-67^+^ hepatocytes and mitotic figures around the peak of proliferation (*i.e*. 48 hrs) after PH, having no effect or increasing these regenerative indices in the consecutive experiments. Hepatic bile salt content, an important determinant of PH-induced liver regeneration, at this time point was not affected by OCA. After pretreatment of mice with FGF19, a reduced expression of ileal bile salt-regulated genes *Fgf15* and *Slc51b* indicating FGF19-mediated repression of bile salt synthesis was seen, but this did not stimulate postresectional liver regeneration in mice.

**Conclusion:** Despite the activation of hepatic and ileal Fxr as shown by induction of target genes, treatment with OCA or FGF19 did not result in accelerated liver regeneration after PH and liver bile salt content was not influenced. We speculate that bile salt homeostasis and endogenous bile salt signaling is already optimal in unaffected livers for proper progression of regeneration after PH. It will be interesting to study the effects of Fxr agonism on liver regeneration after PH, and prevention of PLF in the context of compromised bile salt homeostasis/signaling prior to PH.

## 1. Introduction

Partial hepatectomy is often the preferred curative treatment for hepatobiliary malignancies. However, only 15-20% of patients are eligible for liver resection, mainly due to extensive disease and a predicted insufficient liver remnant [1]. In healthy patients, up to 75% of hepatic volume can be resected [2]. In patients with hepatic functional impairment (*e.g*. due to steatohepatitis or cirrhosis), a maximum of 60% can be removed. An imbalance between liver volume and quality, with lack of functional recovery after (extended) resection, may lead to postresectional liver failure (PLF). PLF is clinically characterized by hyperbilirubinemia, coagulopathy and hepatic encephalopathy, and occurs in up to 9% of patients, with high lethality [1]. Current clinical practice focuses on preoperative enlargement of future remnant liver volume/function (portal vein embolization) [3], or acceleration of functional hypertrophy of the future remnant liver with novel surgical techniques such as the ALPPS (Associating Liver Partition and Portal vein ligation for Staged hepatectomy) procedure [4,5] to overcome this problem. However, both procedures are not always successful, and complication rates are considerable in case of ALPPS.

Postresectional liver regeneration comprises a complex biological response involving interaction between parenchymal and non-parenchymal cells by means of endo-, angio- and paracrine signaling by cytokines, growth factors and metabolic factors [6–8]. In the past decade, bile salts have been recognized as essential signaling molecules in liver regeneration, opening new areas of therapeutic exploration as pharmaceutical modulation of bile salt receptors is evaluated in numerous clinical trials [9,10]. Data from animal models indicate that endogenous (*i.e*. bile salts, BS) or (semi)synthetic agonists (*e.g*. obeticholic acid, OCA) of the ligand-activated transcription factor Farnesoid X Receptor (FXR) can stimulate liver regeneration (LR) in the context of portal vein embolization or after partial hepatectomy (PH) [11–15]. Furthermore, feeding a cholic acid-enriched diet elicits liver growth in the absence of liver resection [11,16].

FXR is highly expressed in the small intestine and liver, but also in the adrenal glands and kidneys [17]. FXR plays a central role in maintaining BS homeostasis and, accordingly, acts to limit detrimental effects (*e.g*. cell death) of BS overload [17,18]. Target genes of FXR include, amongst others, transporters engaged in uptake (*NTCP*) and secretion (*e.g. BSEP, SLC51A/B)* of BS, and genes involved in regulation of bile salt synthesis (*e.g*. intestinal *FGF19/Fgf15* and hepatic small heterodimer partner (*SHP*). Endocrine fibroblast growth factor (FGF) 19/15 and SHP both target, albeit through distinct routes, the *CYP7A1* gene that encodes the rate-limiting enzyme in the dominating BS synthetic pathway [17,19].

BS entry into the ileal enterocyte is the trigger for FXR-mediated transcriptional induction of *FGF19* expression, resulting in enhanced secretion of the enterokine FGF19 (or Fgf15 in rodents) into the portal circulation. The role of Fgf15 in liver regeneration was first demonstrated by Uriarte *et al*. who showed the importance of Fxr and Fgf15 in maintaining BS homeostasis in the regenerating liver remnant.[16] Knocking out either *Fxr* or *Fgf15* led to excessive intrahepatic accumulation of bile salts, increased hepatocellular injury and high mortality in the first 3 days after PH [11,16]. Mouse studies revealed that *Fgf15^-/-^* mice have increased amounts of hepatic Cyp7a1 at mRNA, protein and functional level [20]. Consequently, liver injury and mortality after PH were negated by intraperitoneal adenoviral-mediated *Fgf15* delivery [16]. Moreover, this study also showed that Fgf15 mediates enhanced liver proliferation following BS feeding [16]. Although signaling actions of BS thus seem required for proper postresectional LR, a tight control of intracellular BS levels seems indispensable. In addition, Fgf15 appears to exert a direct mitogenic effect on hepatocytes, since knockdown of its receptor fibroblast growth factor receptor 4 (Fgfr4) impaired hepatocyte proliferation after PH [21]. This was probably due to abrogation of Stat3 signaling which is downstream of Fgfr4, and responsible for induction of *Foxm1b* and subsequent cell cycle progression. In addition to direct mitogenic effects on cultured hepatocytes, FGF19 was reported to enhance growth of cultured cholangiocytes as well [16].

The aim of this study was to investigate the effect of stimulation of the Fxr/Fgf15 pathway on LR after PH in mice. For this purpose, we used the potent FXR agonist OCA, which is approved for treatment of primary biliary cholangitis patients unresponsive to first-line therapy, and is undergoing further clinical evaluation in patients with non-alcoholic steatohepatitis [9].

## 2. Materials and Methods

### 2.1 Animal studies

Male C57BL6/J mice (n=36, 11 weeks old) were obtained from Janvier Labs (Le Genest-Saint-Isle, France) and housed in the animal facility of the Université Catholique de Louvain (UCL; Brussels, Belgium). The animals were kept under controlled conditions with exposure to a 12-h light/12-h dark cycle and a constant temperature of 20-22°C. Animal experiments were conducted in accordance with European regulations and FELASA guidelines for humane care for laboratory animals provided by UCL. The study protocol was approved by the university ethics committee (2012/UCL/MD/026).

### 2.2 PH model

In the first study, mice were fed standard chow (SAFE diets A03; Augry, France), and were pre-treated for one week with FXR agonist obeticholic acid (OCA dissolved in 0.5% methylcellulose; 10 mg/kg, daily oral gavage, n=18) or vehicle (0.5% methylcellulose, n=18). OCA was generously provided by Intercept Pharmaceuticals Inc., New York, USA. After 7 days of pre-treatment, all mice underwent 70% PH (T=0). PH was performed under isoflurane anesthesia essentially according to the protocol of Mitchell and Willenbring [22], with the adaptation that instead of an open abdominal procedure, only a small incision below the xyphoid was made through which the liver was mobilized. After resection of the median and lateral lobes and gallbladder, daily oral gavage with OCA and vehicle continued. BrdU (50 mg/kg) was administered *i.v*. two hours before sacrifice to allow immunohistochemical analysis of hepatic DNA synthesis. At the time of sacrifice (24, 48 or 72 hours after PH, n=6 per treatment group and time point), mice were anesthetized via intraperitoneal injection with ketamine/xylazine. The abdominal cavity was opened via midline incision and blood was drawn by portal vein puncture, kept on ice, and serum was prepared and stored at -80°C until further use. Due to a technical issue, serum was unfortunately not available for biochemical analyses. Part of the liver (‘posterior lobes’) and ileum was fixated in 4% paraformaldehyde, with the remainder of the tissues being snap-frozen in liquid nitrogen and stored at -80°C until analysis.

In the second study, mice were pre-treated for one week with OCA (10 mg/kg, daily oral gavage, n=21) or vehicle (0.5% methylcellulose, n=13). As positive control, an additional group of mice underwent daily intraperitoneal injection with FGF19 (1 mg/kg body weight in sterile PBS, n=13). Recombinant FGF19 was kindly provided by Genentech, San Francisco, USA. All mice underwent PH after 7 days pre-treatment (n=8 or n=16 per group), or were sacrificed (n=5 per group) to obtain baseline measures. Half of the OCA-treated mice received water supplemented with sucrose (42 g/L) *ad libitum*, to prevent post-PH metabolic derangements. Two days after PH, *i.e*. around the peak of hepatocyte proliferation, all mice were sacrificed by exsanguination. Tissue and blood were processed in the aforementioned manner.

#### 2.2.1 Liver mass recovery

The rate of liver mass recovery was estimated using the following formula:

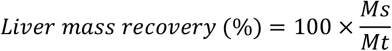

where M_s_ is the liver weight at sacrifice, and Mt is the total liver mass before PH (estimated by dividing the mass of resected segments by 0.7).

### 2.3 Immunohistochemistry

Hepatocyte proliferation was assessed via immunohistochemical staining of Ki-67 and BrdU on serial tissue sections of 4 μm thickness. Mouse monoclonal antibodies against Ki67 (1:50; Code No. M7249, Dako, Glostrup, Denmark) and BrdU (1:100; Code No. M 0744, Dako) were used. Anti-mouse Envision system (Dako) was used for secondary detection. For visualization, the 3,3’-diaminobenzidine (DAB) Substrate-Chromogen System (Dako) was used. Nuclei were counterstained with hematoxylin. The proliferative index (%) was determined by dividing the amount of Ki-67 positive hepatocyte nuclei by the total number of hepatocyte nuclei in five high-power (40x) fields. Mitotic figures were assessed from H&E stained liver sections.

### 2.4 Total bile salt assay

A 5% homogenate of liver tissue was made by homogenizing ca. 50 mg of tissue in 1 mL of 75% ethanol by means of a Mini-Beadbeater (Biospec Products, Bartlesville, USA). Thereafter, BS were extracted by incubating homogenates for 2 hours at 50°C [23]. Supernatant was collected after centrifugation (10 min., 20620*g* at 4°C). The amount of BS present in serum or liver extract was measured via an enzymatic assay (total bile acid assay, Diazyme Laboratories, Dresden, Germany) according to the manufacturer’s protocol. The amount of BS present in liver extract was normalized to liver protein content as measured by bicinchoninic acid (BCA) assay (Pierce^®^ BCA protein assay kit, Thermo Scientific, Waltham, Massachusetts).

### 2.5 Analysis of gene expression

Expression levels of genes involved in BS homeostasis, cell cycle regulation and proliferation, cellular stress response and cytokine regulation were measured in liver and ileum samples via real-time PCR. To study the effect of PH, samples of the quiescent liver (Resected lobes, T=0) were included. RNA was isolated from the resected liver lobe (T=0) and ileal tissue (T=24), and the regenerated lobe at sacrifice (T=48) with TRI reagent solution according to the manufacturer’s protocol (Sigma-Aldrich, St. Louis, Missouri). The concentration and purity of RNA in the samples were determined by measuring absorbance with the NanoDrop 1000A spectrophotometer (Thermo Scientific). After RNA was treated with DNAse (Promega, Madison, Wisconsin), efficiency of DNAse treatment was verified by PCR using primers for an intron-less gene. Next, 750 ng total RNA was reverse transcribed to form cDNA according to the manufacturer’s protocol (SensiFAST™ cDNA Synthesis Kit, Bioline, Luckenwalde, Germany). Real-time PCR analysis was performed according to the manufacturer’s protocol (SensiMix™ SYBR^®^ & Fluorescein Kit, Bioline) with a MyiQ Single-Color Real-Time PCR detection system (Bio-Rad, Veenendaal, the Netherlands). The reaction mixture contained 2.0 μL diluted cDNA sample (corresponding to 7.5 ng total RNA) in a total volume of 10 μL. LinRegPCR software was used to calculate relative expression values [24]. Results were normalized using *Rplp0/36b4* as a reference gene. Data are expressed relative to the median of the control group at baseline (T=0).

### 2.6 Statistical analysis

Data were statistically analyzed using IBM SPSS Statistics version 24 for Microsoft Windows^®^. Non-normal distribution was assumed and groups were compared using the Mann-Whitney U test. In case of more than two groups, the Kruskal-Wallis test was applied. If the Kruskal-Wallis test showed significance (P≤0.05), post-hoc Mann-Whitney U tests were performed within groups with Bonferroni-Holm correction for multiple comparisons. P-values ≤0.05 were considered statistically significant. For visual purposes, data on body weight and glycemia in graphs is depicted as mean ± SEM. All experimental data on gene expression and biochemical analyses are graphically presented as median with interquartile range.

## 3. Results

### 3.1 OCA pre-treatment modulates intestinal and hepatic Fxr target gene expression

To test the hypothesis that FXR activation augments LR after PH, we examined the effect of OCA on liver regeneration at 24, 48 and 72 hours after PH (**Fig. 1**). Prior to PH, mice received daily OCA gavage for 7 days.

**Figure 1.**
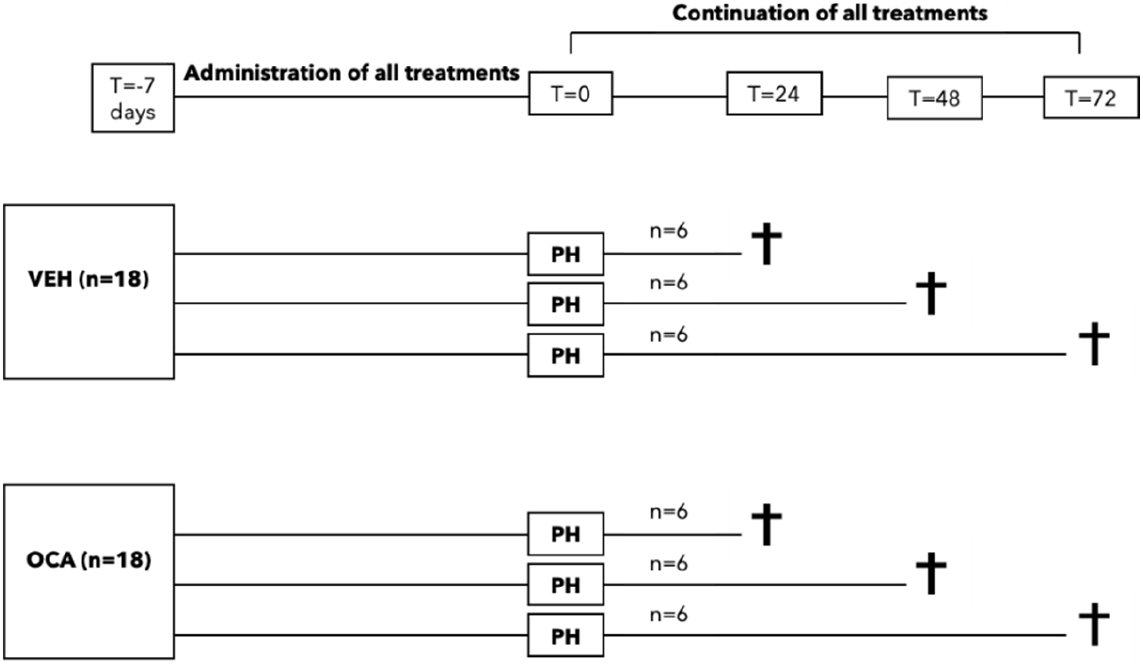
Study design. Mice (n=6 mice per group) were pre-treated for 7 days by daily administration of OCA (10 mg/kg/day) or vehicle, before undergoing 70% PH (T=0) with continuation of treatments. Mice were sacrificed 24, 48 and 72 hours later. VEH, vehicle; OCA, obeticholic acid; PH, partial hepatectomy.

Effectiveness of OCA pre-treatment was inferred from elevated ileal expression of Fxr target genes *Fgf15* and *Slc51b* (p<0.010) (**Fig. 2AB**). Additional Fxr target genes were also modulated, *viz. Cyp8b1* (-3.3 fold; *p=*0.002) and *Bsep* (*p=*0.004) in the liver (**Fig. 2CD**). Expression of hepatic *Fxr* per se was not affected by OCA treatment (**Fig. 2E**).

**Figure 2.**
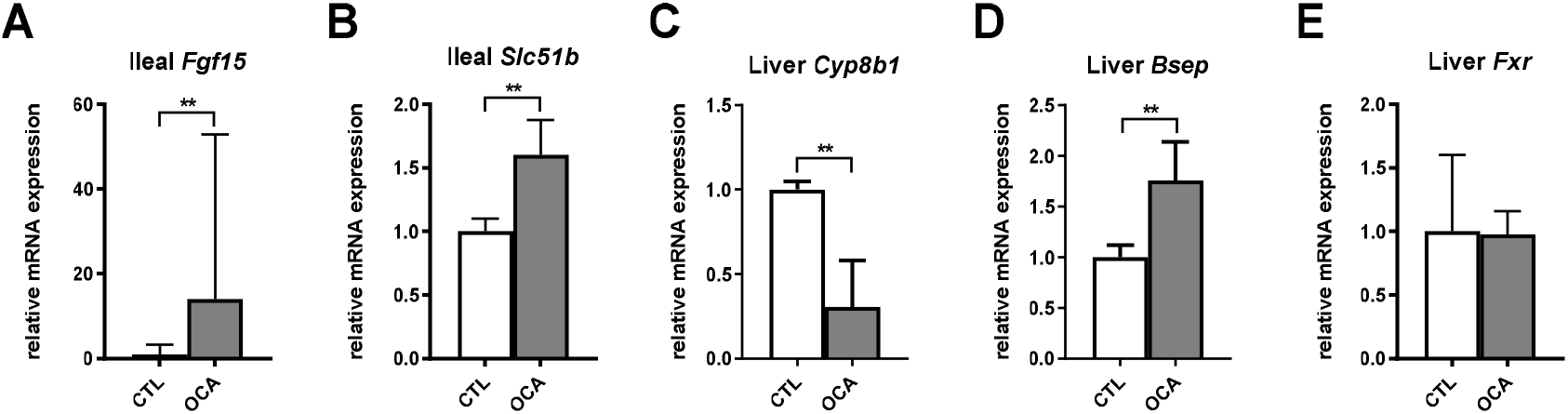
Effect of pre-treatment with OCA (10 mg/kg/day) on expression of Fxr target genes. Mice (n=6 mice per group) were pre-treated for 7 days by daily administration of OCA or vehicle, before undergoing 70% PH. Transcripts were analyzed in the terminal ileum of mice sacrificed 24 hrs after PH (panels **AB**), and in the liver (segments resected at T=0, panel **C-E**). Values are expressed relative to the median expression in the control group, and displayed as median with interquartile range. **p<0.01. OCA, obeticholic acid; *Fgf*15, fibroblast growth factor 15; *Slc51b*, organic solute transporter 51 beta; *Cyp8b1*, sterol 12α-hydroxylase; *Bsep*, bile salt export pump; *Fxr*, farnesoid x receptor.

#### 3.1.1 OCA has no effect on functional liver generation parameters

OCA had no effect on body weight (**Fig. 3A**) or glycaemia course (data not shown) during the pre-treatment period. PH resulted in a transient drop in body weight in the first 2 days after surgery, with body weight returning to pre-surgical values at T=72h in the control group. Of note, OCA-treated mice continued to lose weight after post-operative day 2.

**Figure 3.**
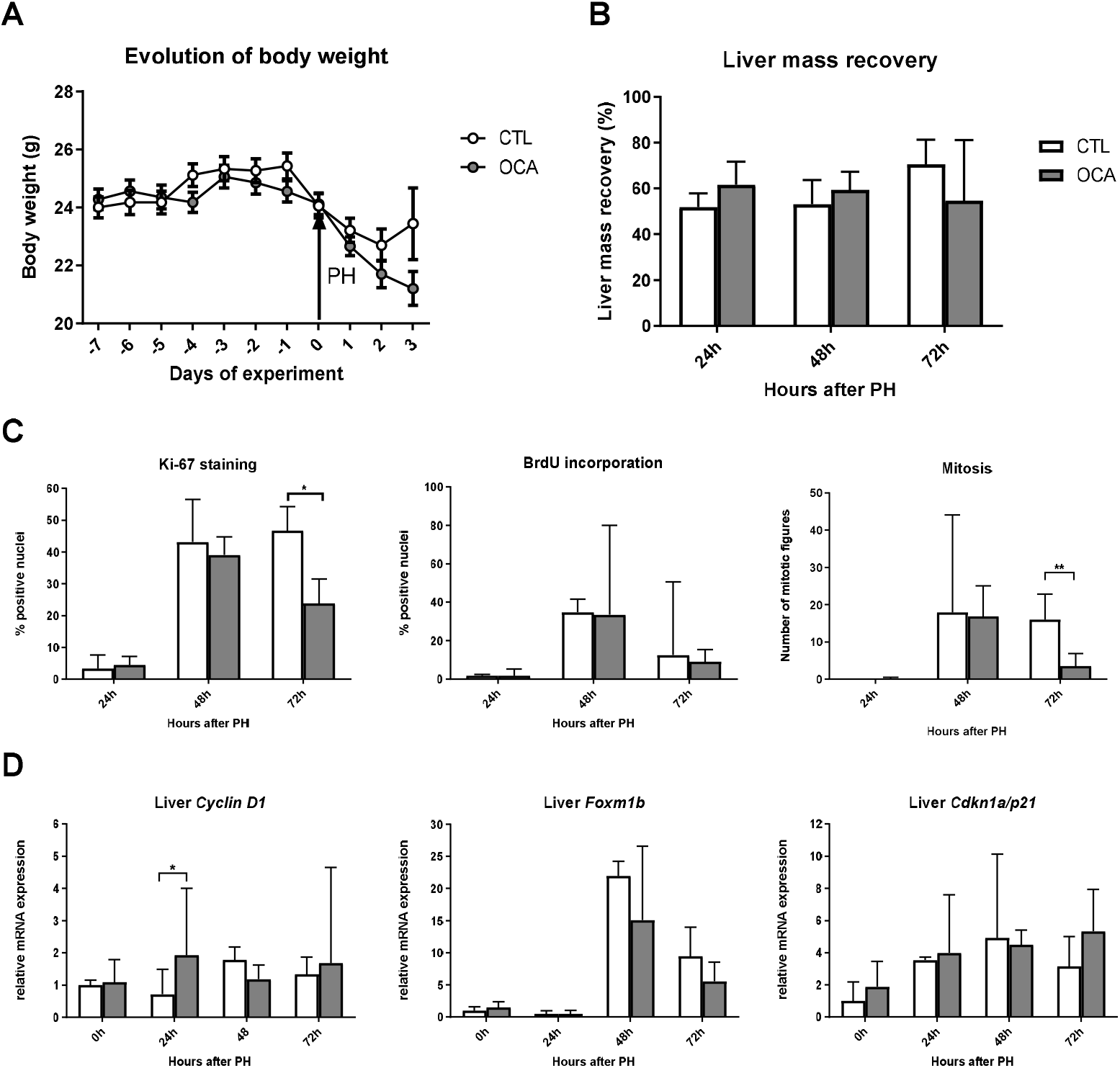
Effect of pre-and posttreatment with OCA on functional parameters, proliferative measures and mitotic gene expression in the liver. Mice (n=6 mice per group) were pre-treated for 7 days by daily administration of OCA or vehicle, before undergoing 70% PH. Mice were sacrificed at 24, 48 and 72 hours after PH. Body weight (**A**) was recorded daily from 7 days before surgery till sacrifice. Regeneration after PH was assessed by recovery of liver mass (**B**) and immunohistochemical/histological analysis of hepatocyte proliferation (**C**). Transcripts were analyzed in the liver (**D**), and values are expressed relative to the median expression in the control group. Data are depicted as median with interquartile range except for data on body weight evolution (mean ± SEM) *p<0.05. **p<0.01. OCA, obeticholic acid; PH, partial hepatectomy; BrdU, bromodeoxyuridine; Fox, forkhead box protein; Cdkn, cyclin-dependent kinase inhibitor.

Liver mass recovery was similar in OCA-treated animals and controls at each of the studied time points after PH (**Fig. 3B**). DNA synthesis has been reported to increase at 32 hours post-hepatectomy in mice, with an initial peak reflecting hepatocyte division at 36-40 hours [25]. Here, we observed a peak in nuclear Ki67 staining in hepatocytes, BrdU incorporation and mitotic events at 48 hours after PH (**Fig. 3C**). OCA treatment did not result in an earlier peak and had no effect on the percentage of Ki-67^+^ hepatocytes, BrdU incorporation or the number of mitotic figures at T=48h. In fact, the number of mitotic figures and Ki-67^+^ nuclei at T=72h was decreased in OCA-treated mice (both p<0.05). Above observations are not in support of our hypothesis. Note that there was no sign of histological injury, cholestasis or necrosis, at each of the respective points of sacrifice, in any of the groups (data not shown).

Next, we determined whether PH elicited expected changes in expression of key genes in cell cycle progression. Cyclin D1, pivotal for progression of hepatocytes through G1 phase, starts increasing at approximately 30 hours after PH (prior to peak DNA synthesis) [26–28]. Expression of *CyclinD1* was increased in the OCA-treated animals at 24 hours after PH in comparison to the control group (**Fig. 3D**). This did not translate into functional changes later in the cell cycle or regenerative course (*i.e*. liver mass recovery, number of mitotic figures, or percentage of Ki-67^+^ hepatocytes). PH resulted in strong induction of *Foxm1b* from T=48h onwards. The extent of upregulation was similar in both groups at T=48h and T=72h, despite *Foxm1b* being an Fxr target gene [29]. Liver expression of *Cdkn1a/p21*, involved in G1 phase cell cycle arrest, was upregulated in both groups after PH.

#### 3.1.2 OCA reduces BS levels in the regenerating liver

Maintenance of BS homeostasis is essential for normal progression of liver regeneration after PH [16]. To see whether lack of effects of OCA on liver regrowth was due to disturbed BS homeostasis, we determined hepatic BS content in the quiescent (resected lobes) and regenerating liver. OCA led to decreased BS content in the remnant liver, which reached significance at T=24h (**Fig. 4A**). Note that there was substantial within-group variation in hepatic bile salt content in control mice following PH. Marked repression of *Cyp7a1* was evident after PH, with superimposed downregulation by OCA at all time points after PH (**Fig. 4B**). In addition to reduced BS synthesis, enhanced basolateral efflux of BS via upregulated expression of *Slc51b* may have contributed to reduced BS content in the OCA-treated group (**Fig. 4C**). Bsep was upregulated by OCA solely before PH (**Fig. 4D**).

**Figure 4.**
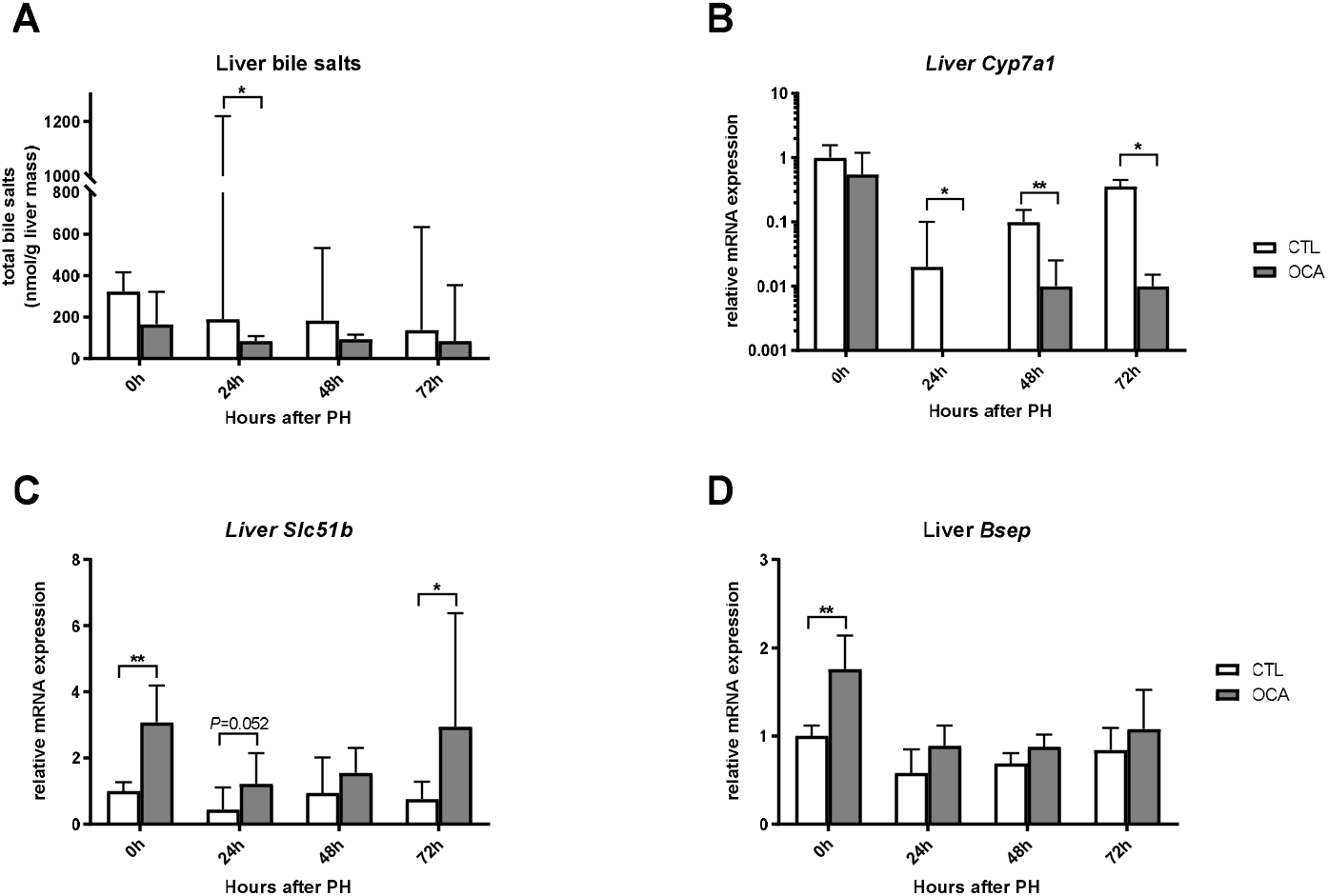
Effect of pre- and post-treatment with OCA on hepatic bile salt levels and Fxr-regulated pathways in the liver. Mice (n=6 mice per group) were pre-treated for 7 days by daily administration of OCA or vehicle, before undergoing 70% PH, with continuation of treatments thereafter. Mice were sacrificed at 24, 48 and 72 hours after PH. Total bile salts levels in the liver (**A**) were analyzed. Transcripts (**B-D**) were analyzed in the liver, and values are expressed relative to the median expression in the control group. Data are presented as median with interquartile range.*p<0.05. **p<0.01. OCA, obeticholic acid; PH, partial hepatectomy; *Cyp7a1*, cholesterol 7α-hydroxylase; *Slc51b*: organic solute transporter 51 beta; *Bsep*, bile salt export pump.

Post-PH metabolic derangements (no body weight recovery, reduced serum glucose) and poorer well-being (scored by, amongst others, rough hair coat, squinted eyes, hunched walking) were observed in OCA-treated mice, and this may have masked effects of OCA on liver regeneration after PH. Therefore, in a second experiment, half of the group of hepatectomized mice given OCA, received water supplemented with sucrose. To see whether liver regeneration could be enhanced under our experimental conditions, an additional group of mice was treated with recombinant FGF19, which was previously shown to stimulate hepatocellular proliferation [16]. Five mice per group were sacrificed after pre-treatment for one week without undergoing PH, to determine baseline effects of OCA and FGF19 pre-treatment. The other mice were subjected to 70% PH and sacrificed 48 hrs later (**Fig. 5**).

**Figure 5.**
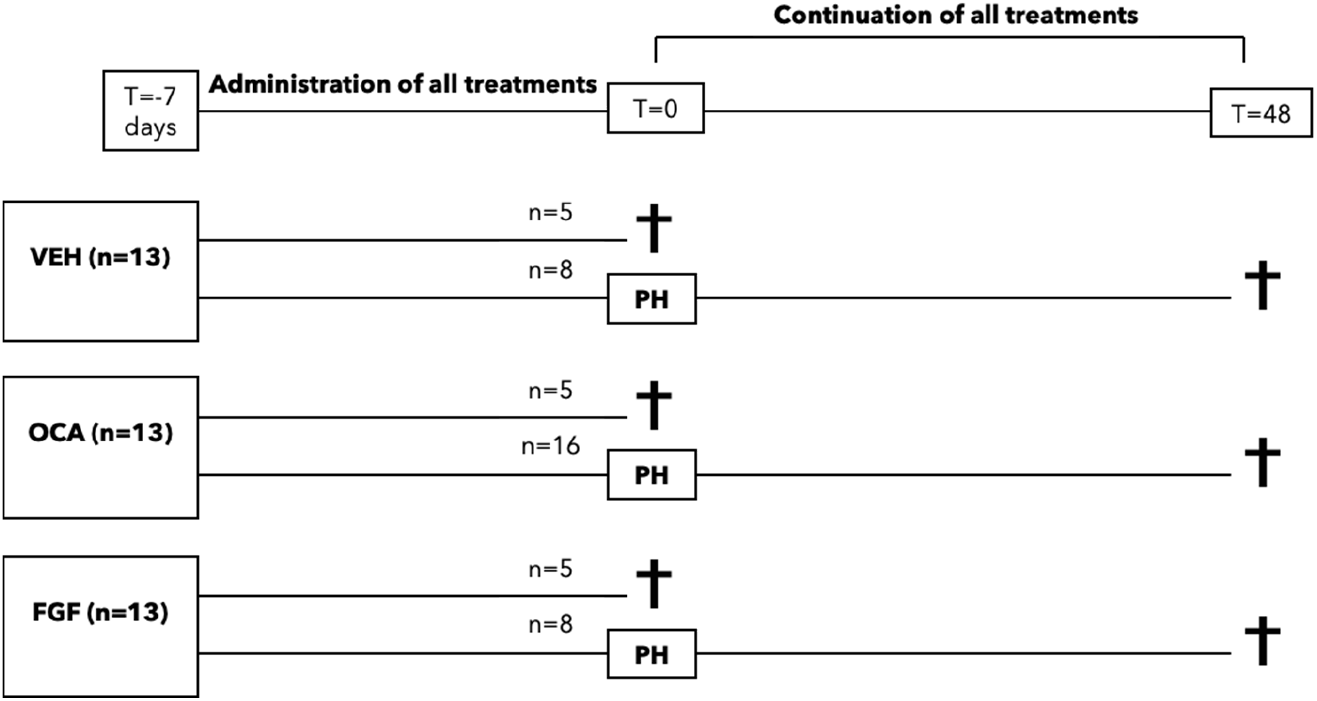
Study design of the follow-up experiment. Mice were pre-treated for 7 days by daily administration of OCA, FGF19 or vehicle, before sacrifice (n=5) or 70% PH with continuation of initial treatments (n=8). Mice that were subjected to 70% PH were sacrificed 48 hrs later. VEH, vehicle; OCA, obeticholic acid; FGF, fibroblast growth factor 19; PH, partial hepatectomy.

### 3.2 Effects of pre-treatment with OCA and FGF19

Effects of pre-treatments with OCA and FGF19 were assessed in mice sacrificed without undergoing PH. OCA treatment resulted in downregulation of hepatic *Cyp8b1* (**Fig. 6A**), but had no significant effect on *Cyp7a1* **(Fig. 6B)**. We confirmed an unchanged hepatic *Fxr* expression after Fxr activation (**Fig. 6C**) [30]. Unexpectedly, and in contrast to the first experiment, OCA pre-treatment had no effect on liver expression of *Bsep* (**Fig. 6D**) and *Slc51b* (data not shown). This is not readily explained. In the first experiment, baseline effects were derived from resected segments with potential superimposed effects of liver mobilization and surgical manipulation [31], but this is unlikely to explain the discrepancy. OCA tended to induce expression of ileal *Fgf15* (post-hoc test *p=*0.151), whilst *Slc51b* expression was unaltered (**Fig. 6EF**). As discussed later, transcriptional effects of OCA appeared more prominent in the post-hepatectomy phase **(Fig. 9CD).**

**Figure 6.**
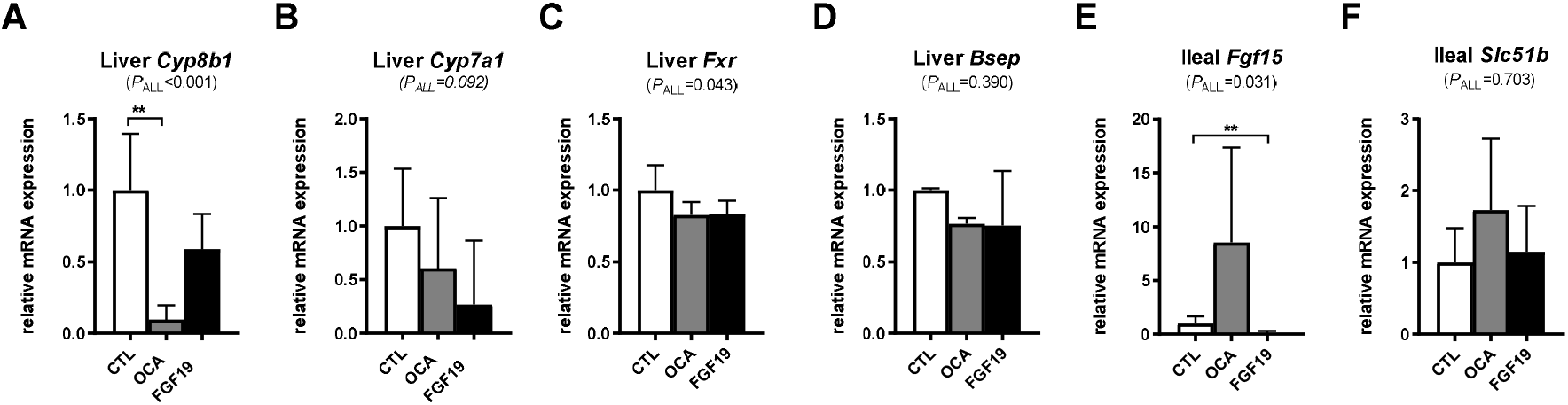
Effect of pre-treatment with OCA or FGF19 on Fxr/FGF19-regulated pathways in the liver and ileum. Mice (n=5 mice per group) were pre-treated for 7 days by daily administration of OCA, FGF19 or vehicle, before sacrifice. Transcripts were analyzed in the liver (**A-D**), and the terminal ileum (**E-F**). Values are expressed relative to the median expression in the control group. Data are displayed as median with interquartile range. **p<0.01 (post-hoc test). CTL, vehicle-treated mice, OCA, obeticholic acid; FGF, fibroblast growth factor; *Cyp8b1*, sterol 12α-hydroxylase; *Cyp7a1*, cholesterol 7α-hydroxylase; *Fxr*, farnesoid X receptor; *Bsep*, bile salt export pump; *Slc51b*,organic solute transporter 51 beta.

Effectiveness of FGF19 treatment was tested by studying expression of *Cyp7a1*, which is repressed as consequence of binding of FGF19 to its hepatic receptor Fgfr4. Unexpectedly, *Cyp7a1* levels showed a trend but were not significantly affected by FGF19 treatment (**Fig. 6B**). In contrast, ileal expression of the bile salt-regulated gene *Fgf15* was reduced by FGF19 administration (**Fig. 6E**), which can be interpreted as a secondary consequence of FGF19-mediated repression of bile salt synthesis, with a smaller supply of bile salts reaching the small intestine. FGF19 did not affect hepatic expression of *Cyp8b1* and *Bsep* (**Fig. 6AD**). Hepatocyte receptors *Fgfr4* and *Klb*, for which FGF19/Fgf15 is a ligand, were expressed to the same extent in all groups (**Fig. S1**).

We next determined effects of OCA and FGF19 on liver regeneration at 48 hrs after PH. Mice treated with OCA and receiving sucrose-supplemented water in the post-PH course, were indistinguishable on all examined parameters (body weight, glucose, well-being, serum biochemistry, bile salt levels, gene expression) from OCA-treated mice receiving plain water (data not shown). We therefore decided to merge data from these groups into a single group of n=16 (**Fig. 7AB**). Levels of circulating liver enzymes and bilirubin at 48 hrs after resection were similar among groups (**Fig. 7C**), showing a 1.5-2.0-fold increase in all groups compared to baseline values (data not shown). Liver mass recovery after PH was comparable among groups (**Fig. 7D**), whilst the number of Ki-67^+^ hepatocyte nuclei and mitotic figures were significantly increased (p<0.001) in the OCA-treated mice (**Fig. 7D**).

**Figure 7.**
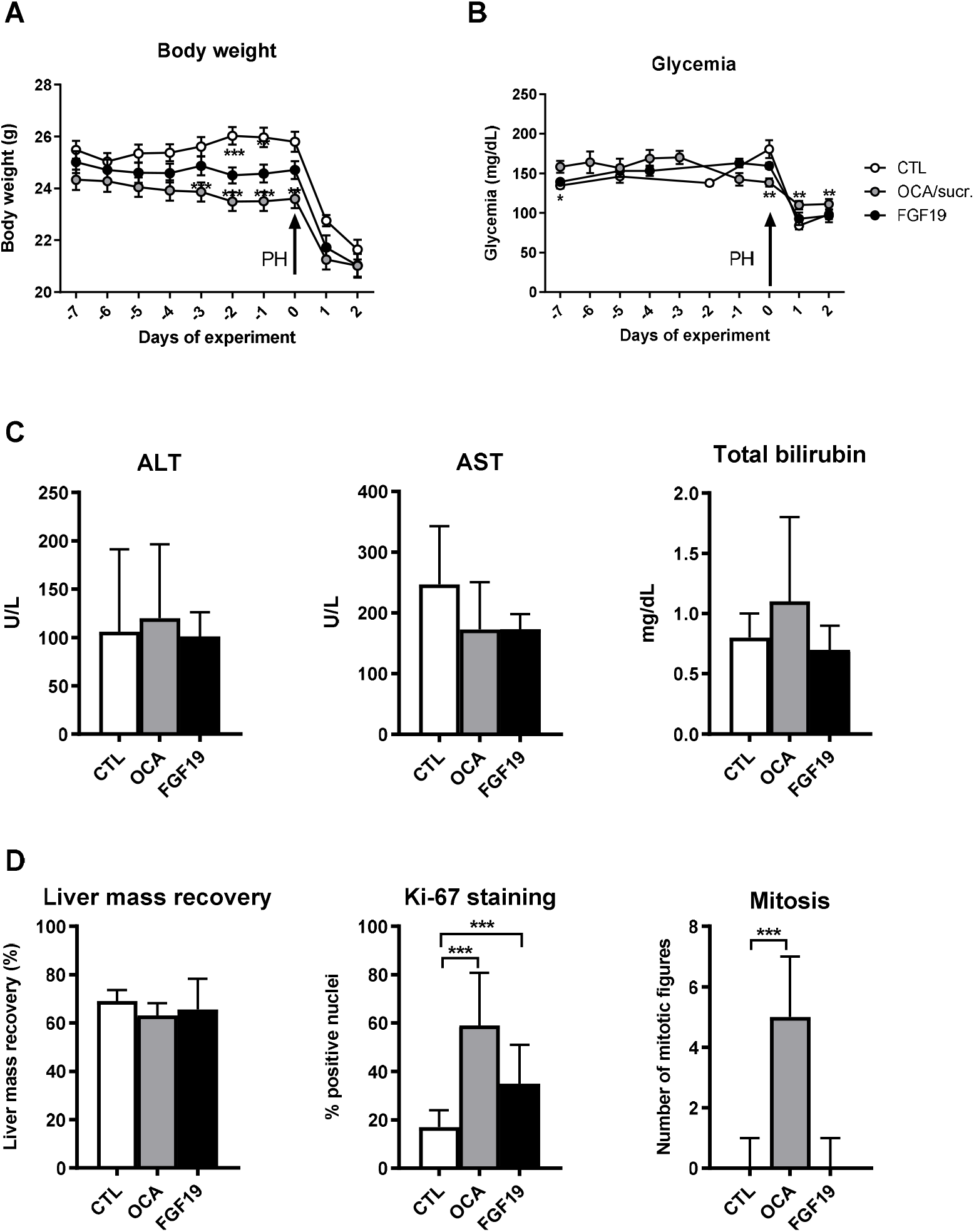
Effect of treatment with OCA or FGF19 on functional parameters, liver biochemistry and proliferative measures after partial hepatectomy. Mice (n=8 or 16 mice per group) were pre-treated for 7 days by daily administration of OCA, FGF19 or vehicle, before undergoing 70% PH and continuation of treatments. Half of the group of OCA-treated mice received sucrose-supplemented water in the post-hepatectomy phase, data from the entire group receiving OCA was analyzed collectively as explained in the text. Mice were sacrificed at 48 hours after PH. Body weight (**A**) and glycemia levels (**B**) were recorded daily from 7 days before surgery to sacrifice. Liver enzymes and total bilirubin were determined in serum at exsanguination (**C**). Regeneration after PH was assessed by recovery of liver mass and immunohistochemical/histological analysis of hepatocyte proliferation (**D**). Data are depicted as median with interquartile range except for data on body weight and glucose evolution (mean ± SEM) **p<0.01. ***p<0.001. CTL, vehicle-treated mice; OCA/sucr., obeticholic acid-treated mice, with half of the group receiving post-operative sucrose supplementation; FGF19, fibroblast growth factor 19; PH, partial hepatectomy; ALT, alanine aminotransferase; AST, aspartate aminotransferase.

#### 3.2.1 Effect of OCA and FGF19 on cell cycle progression after PH

Increased proliferation after OCA treatment was not corroborated by upregulation of Fxr target gene *Foxm1b* (**Fig. 8A**) and its downstream target gene *Cdc25b* (**Fig. 8B**) [11]. Expression of cyclins *Ccnd1* and *Ccne1* was not increased compared to control animals (**Fig. 8CD**). In contrast, a significant upregulation of pivotal regulator of cell cycle progression *Ccna2* was detected in mice treated with OCA (**Fig. 8E**).

**Figure 8.**
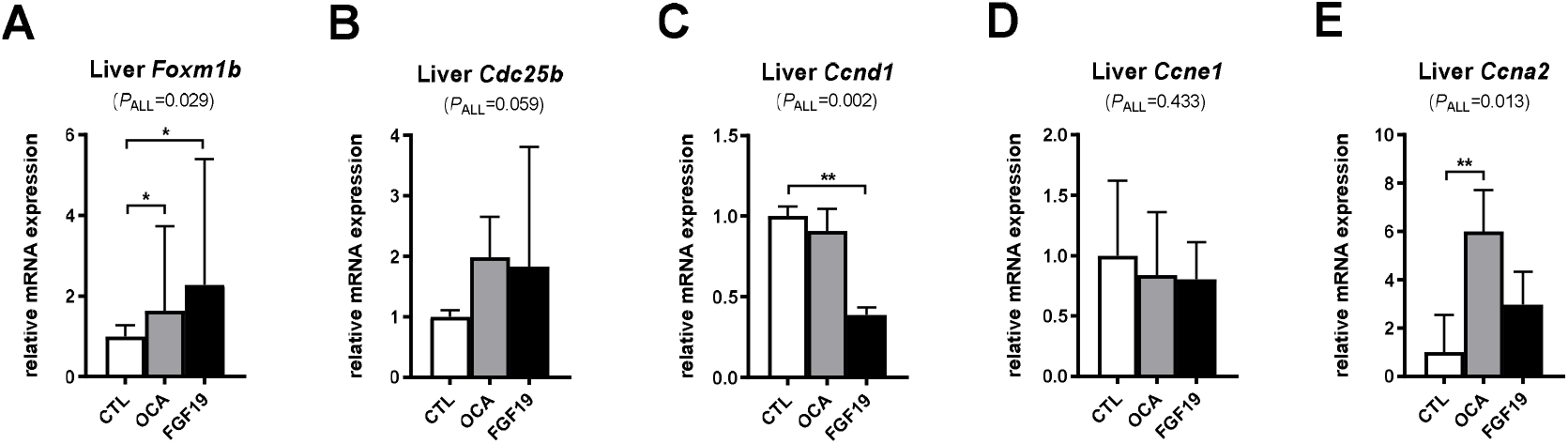
Effect of treatment with OCA or FGF19 on expression of cell cycle regulatory genes in hepatectomized mice. Animals (n=8 or 16 mice per group) were pre-treated for 7 days by daily administration of OCA, FGF19 or vehicle, before undergoing 70% PH and continuation of treatments. Mice were sacrificed at 48 hours after PH. Transcripts (**A-E**) were analyzed in the liver (segments resected at T=48). Values are expressed relative to the median expression in the control group. Data are presented as median with interquartile range. **p<0.01. CTL, vehicle-treated mice; OCA, obeticholic acid; FGF19, fibroblast growth factor 19; *Fox*, forkhead box protein; *Ccn*, cyclin.

In animals treated with FGF19, *Foxm1b* did not change after PH, whilst expression of *Ccnd1* was decreased (**Fig. 8AC**). No significant upregulation of cell cycle progression regulator *Cdc25b* was seen compared to the control group, which was similar for *Ccne1* and *Ccna2* (**Fig. 8BDE**).

Serum and hepatic bile salt content was not different among groups (**Fig. 9AB**), despite a reduced expression of *Cyp7a1* in mice treated with OCA (**Fig. 9C**). A decrease in *Cyp7a1* expression was not seen in the animals receiving FGF19, suggesting that co-activation of the ileal and hepatic axis is important for optimal repression of *Cyp7a1*. The time interval between administration of FGF19 and liver harvesting might also be a reason for the lack of effect of FGF19 on *Cyp7a1* expression. Mice were sacrificed 0.5-2.5 hrs after the last administration of OCA/FGF19, whilst studies in HepG2 hepatoma cells indicate that it takes at least 2 hrs before *CYP7A1* mRNA is significantly reduced by FGF19 (data not shown). Expression of *Ntcp* was also significantly reduced in mice receiving OCA (**Fig. 9D**).

**Figure 9.**
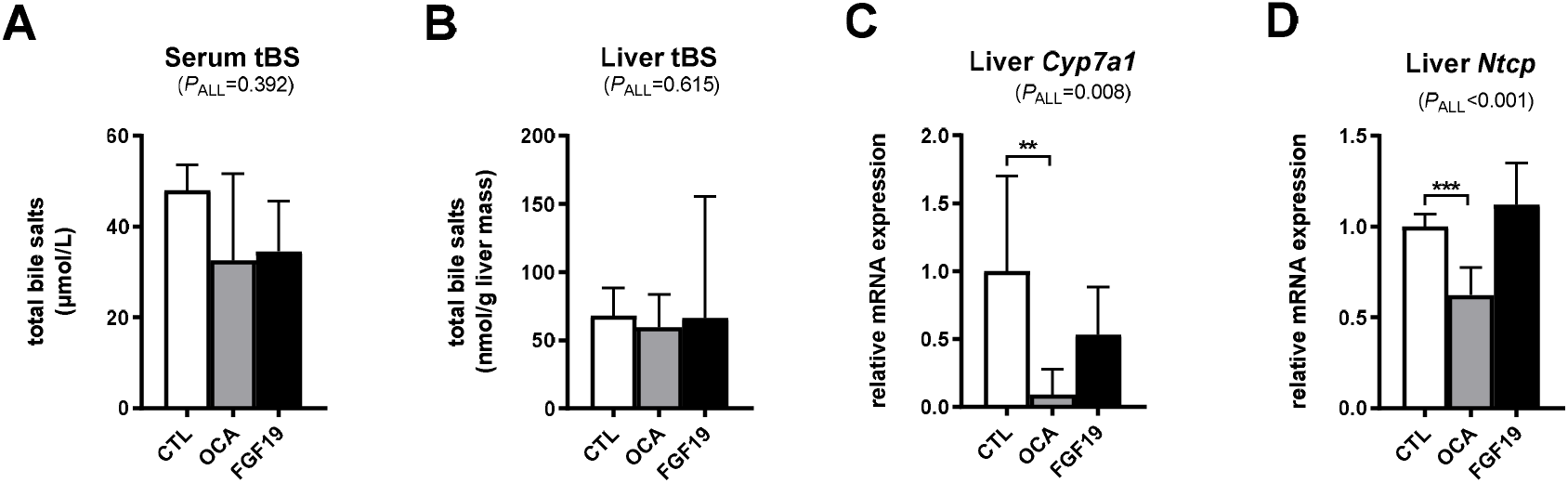
Effect of treatment with OCA or FGF19 on bile salt levels and genes involved in bile salt synthesis and uptake in hepatectomized mice. Animals (n=8 or 16 mice per group) were pre-treated for 7 days by daily administration of OCA, FGF19 or vehicle, before undergoing 70% PH and continuation of treatments. Mice were sacrificed at 48 hours after PH. Total levels of serum and liver bile salts **(A-B)** and hepatic transcripts **(C-D)** were analyzed. Values are expressed relative to the median expression in the control group. Data are displayed as median with interquartile range. **p<0.01. ***p<0.001. CTL, vehicle-treated mice; OCA, obeticholic acid; FGF19, fibroblast growth factor 19; tBS, total bile salts; *Cyp7a1*, cholesterol 7α-hydroxylase; *Ntcp*, sodium-taurocholate co-transporting polypeptide.

## 4. Discussion

Bile salts have emerged as essential signaling molecules in liver regeneration after PH. Data from animal experiments indicate that endogenous bile salts can stimulate liver regeneration and prevent liver injury after PH via the hepatic Fxr and ileal Fxr-Fgf15 axis. Our aim was to investigate whether exogenous activation of the Fxr pathway by the semi-synthetic bile acid derivative OCA (a.k.a. Ocaliva^®^) could stimulate postresectional liver regeneration in mice.

Although modulation of *Cyp7a1, Cyp8b1, Fgf15* and other Fxr targets implied general effectiveness of OCA treatment, no effect of OCA could be detected on liver mass recovery after PH (**Fig. 3B, Fig. 7D**). We observed inconsistent effects of OCA on other regenerative indices in hepatectomized mice, specifically the number of Ki-67^+^ hepatocytes and mitotic figures. OCA had no effect on these indices at 48 hrs after PH in the first experiment (**Fig. 3C),** whereas treatment with OCA resulted in an increased percentage of Ki-67^+^ hepatocytes and number of mitotic figures in the second experiment (**Fig. 7D**). The latter was accompanied by 6-fold higher hepatic expression of cell cycle regulator *Ccna2* **(Fig. 8E).** Hepatic bile salt content, an important determinant of PH-induced liver regeneration [16], was not affected by OCA around the peak of hepatocyte proliferation at 48 hrs after PH (**Fig. 4A, Fig. 9B**). Welfare of the animals was decreased in OCA-treated mice after PH in the first experiment. Taken together, OCA had no consistent effects on regenerative indices other than liver mass recovery, which was indistinguishable in control and OCA-treated mice. Although liver mass regrowth is not an indicator for functional liver recovery *per se*, it is one of the most used parameters to capture this complex cascade of events. This suggests that liver regeneration was already progressing optimally after vehicle treatment and Fxr agonism did not further benefit liver regeneration after 70% PH. It must be noted, though, that liver mass estimation can be influenced by surgical procedure, inter-surgeon variability and amongst others hepatic oedema.

Note that in the first experiment, liver regeneration was studied at multiple time points after PH, with %Ki-67^+^ hepatocytes, BrdU incorporation and number of mitotic figures, peaking in control mice at 48 hrs after PH (**Fig. 3**). For the second experiment only this peak time point was studied, and it appeared that %Ki-67^+^ hepatocytes and especially number of mitotic figures were notably lower (at most 1 mitotic figure per 40x field) in control mice in comparison with the first experiment. Mitotic figures were readily observed until at least the last point of sacrifice (*i.e*. 72 hrs after PH) in the first experiment. With regard to hepatic bile salt content at 48 hrs after PH, absolute levels were dissimilar in control mice of the first and second experiment. The reasons for these discrepancies are unclear.

In the present experiments we studied mice with uncompromised liver function. Compromised livers such as cirrhotic or cholestatic livers, and extended liver resection, are important factors for leaving patients ineligible for PH in clinical practice. Moreover, upon surgery incidences of PLF and mortality are increased in these patient groups. After PH, it is suggested that a relative hepatic overload of potentially toxic bile salts is one of the causative factors for PLF [16]. Cholestasis decreases hepatic regenerative capacity.[32] Comparable rodent models with bile salt overload and decreased liver regeneration comprise an extensive (90%) hepatectomy [16] and bile duct ligation with consequent cholestasis [33,34]. Rodents receiving cholestyramine or pre-operative bile diversion to deplete the whole-body bile salt pool, also showed decreased liver regrowth after PH, emphasizing the necessity of maintaining bile salt signaling and/or homeostasis [11].

A beneficial effect of Fxr agonism on liver regeneration was previously shown by Chen *et al*. who demonstrated that Fxr agonism alleviated the age-related liver regeneration defect in mice[29], highlighting FXR as a potential target for promoting liver regeneration in older patients. Moreover, OCA treatment may increase the efficacy of portal vein embolization and, thereby, resectability. Earlier work by our group has shown that OCA accelerates liver regeneration after experimental portal vein embolization, in terms of liver volume, liver function, and hepatocyte proliferation [12].

The beneficial metabolic effects of FXR agonists, OCA in particular, and FGF19 have been widely studied in many human and experimental settings. Substantial numbers of selective FXR agonists have been tested in rodents to study proliferative and metabolic responses [35,36]. Improved serum enzymes and reduced steatosis after PH were described upon treatment with synthetic FXR agonist GW4064, but only limited proliferative effects were seen [11,37]. Perioperative oral gavage with alisol B 23-acetate (AB23A) resulted in upregulation of Fxr-dependent proliferative genes, amelioration of liver injury, and decreased bile salt synthesis and hepatic bile salt content, after 70% PH in a non-compromised mouse model [38]. Earlier pre-clinical studies demonstrated that OCA treatment results in improvement of non-alcoholic steatohepatitis (NASH) and cirrhosis at the histological level (fibrosis, hepatocellular ballooning, steatosis, and lobular inflammation) [9,39,40], decreased portal hypertension [41], increased metabolic rate [42], and reduced atherosclerosis by increasing fecal cholesterol excretion [43]. Recent studies focusing on intestinal benefits described preservation of intestinal mucosal wall integrity, attenuation of intestinal inflammation and reduced bacterial translocation after OCA treatment in rat models of cholestasis and intestinal ischemia-reperfusion [44,45], and in NASH models [46]. Transient portal hypertension after PH is associated with gut leakiness and portal endotoxemia, the latter contributing to triggering of the regenerative cascade by stimulating macrophage release of pro-inflammatory cytokines [6,47].

For the present study, it cannot be ruled out that improved gut barrier function with concomitant lower levels of LPS in the portal circulation may have masked an enhancing effect of OCA on liver regrowth.

We observed decreased serum glucose levels before and after PH, and significant weight loss after treatment with OCA (**Fig. 7**). These metabolic effects were comparable to data in rodent and human studies [9] and seemed to be accompanied by a decreased score on our animal welfare assessment. It appeared that in the perioperative setting, mice were too vulnerable to cope with these metabolic changes which may have affected liver regrowth as well. Metabolic (side) effects of OCA may be due to the large number of Fxr target genes, many of which remain to be fully characterized [48].

In the second experiment, we added an experimental group that underwent intraperitoneal injection of FGF19 as in the study of Uriarte *et al*. adenoviral-mediated expression of FGF19 abrogated the diminished liver regeneration in *Fgf15^-/-^* mice [16]. Moreover, in livers with a compromised background (*viz*. impaired liver regeneration due to acetaminophen poisoning and partial hepatectomy in aged mice), Alvarez-Sola *et al*. showed that the chimeric FGF19/apolipoprotein A-I molecule Fibapo attenuates liver injury, boosts regeneration as seen by potentiated cell growth-related pathways and increased functional liver mass, and improves survival [49]. Additionally, Fibapo reduces liver bile salt and lipid accumulation, and results in improved fatty liver regeneration and increased survival [50]. However, in our study treatment with FGF19 did not stimulate postresectional liver regeneration. Although reduction of expression of ileal bile salt-regulated genes *Fgf15* and *Slc51b*, a possible consequence of FGF19-mediated repression of bile salt synthesis, was seen after pre-treatment, FGF19 did not affect hepatic expression of bile salt-regulated genes *Cyp7a1, Cyp8b1* and *Bsep* (**Figs. 6 and 9**). After PH, *Foxm1b* was significantly upregulated by FGF19 but without a stimulatory effect on other regulators of cell cycle progression, *viz*. *Cdc25b, Ccne1, Ccna2* and *Ccnd1* (**Fig. 8**). In addition, liver mass recovery or mitotic figures at 48 hrs after PH (**Fig. 7D**) were not affected by treatment with FGF19, although the number of Ki67^+^ hepatocytes was increased. Hepatic bile salt content was not affected by FGF19 treatment around the peak of proliferation at 48 hrs after PH (**Fig. 9B**). Above findings may indicate that the treatment with FGF19 used in the present study, was suboptimal.

Csanaky *et al*. demonstrated that after PH, hepatocytes were protected from bile salt toxicity by increased canalicular and basolateral bile salt secretion resulting in increased serum bile salt levels, whereas total hepatic bile salt content tended to increase but was not significantly influenced by PH [51]. Moreover, a change in hepatic bile salt composition in favor of unconjugated bile salts was observed [51]. Serum bile salt levels were reported to remain elevated until 72 hrs after PH in mice [52]. Conversely, we observed unaltered serum bile salt levels at 48 hrs after partial hepatectomy in all groups but the FGF19-treated mice which had somewhat higher -albeit not significantly different from controls and OCA-treated mice- levels at baseline **(Fig.S2)**. Furthermore, despite upregulation of *Slc51b* and repression of *Cyp7a1* and *Cyp8b1*, hepatic bile salt content did not decrease in OCA-treated animals. In addition, since serum bile salt levels and hepatic bile salt content were not influenced by treatment with OCA prior to and after PH, It would be interesting to examine the effect of administering OCA in the post-PH phase only. In such a set-up, the different groups of mice would share the same metabolic starting point. We speculate that bile salt homeostasis is already optimally maintained for proper progression of liver regeneration after PH. However, in the experimental or clinical setting of compromised bile salt homeostasis/signaling prior to partial hepatectomy, *e.g*. due to external bile diversion or cholestasis, FXR agonism may be of benefit.

Collectively, our results show that OCA modulates expression of hepatic and ileal Fxr target genes. In the first experiment, comprising analyses at 24, 48 and 72 hrs after PH, no effects of OCA on various regenerative indices (liver mass recovery, BrdU incorporation, percentage of Ki-67^+^ hepatocytes, number of mitotic figures) were observed. In the second experiment, comprising analyses at 48 hrs after PH only, OCA treatment gave rise to a higher percentage of Ki-67^+^ hepatocytes and greater number of mitotic figures, but had no effect on liver mass regrowth. We conclude that OCA does not accelerate liver mass recovery after PH in healthy mice.

## Abbreviations

ALT: alanine aminotransferase
ALPPS: Associating Liver Partition and Portal vein ligation for Staged hepatectomy
AST: aspartate aminotransferase
BrdU: bromodeoxyuridine
BS: bile salt(s)
BSEP: bile salt export pump
CCN: cyclin
CDKN: cyclin-dependent kinase inhibitor
CYP7A1: cholesterol 7α-hydroxylase
FGF: fibroblast growth factor
FGFR: fibroblast growth factor receptor
FOX: forkhead box protein
FXR: farnesoid x receptor
LR: liver regeneration
NTCP: sodium-taurocholate co-transporting polypeptide
OCA: obeticholic acid
PH: partial hepatectomy
PLF: postresectional liver failure
SHP: small heterodimer partner
TBS: total bile salts
VEH: vehicle

## Acknowledgements

We are grateful to Intercept Pharmaceuticals for providing obeticholic acid, and Genentech for providing recombinant human FGF19.

## Supplemental Information

**Figure S1.**
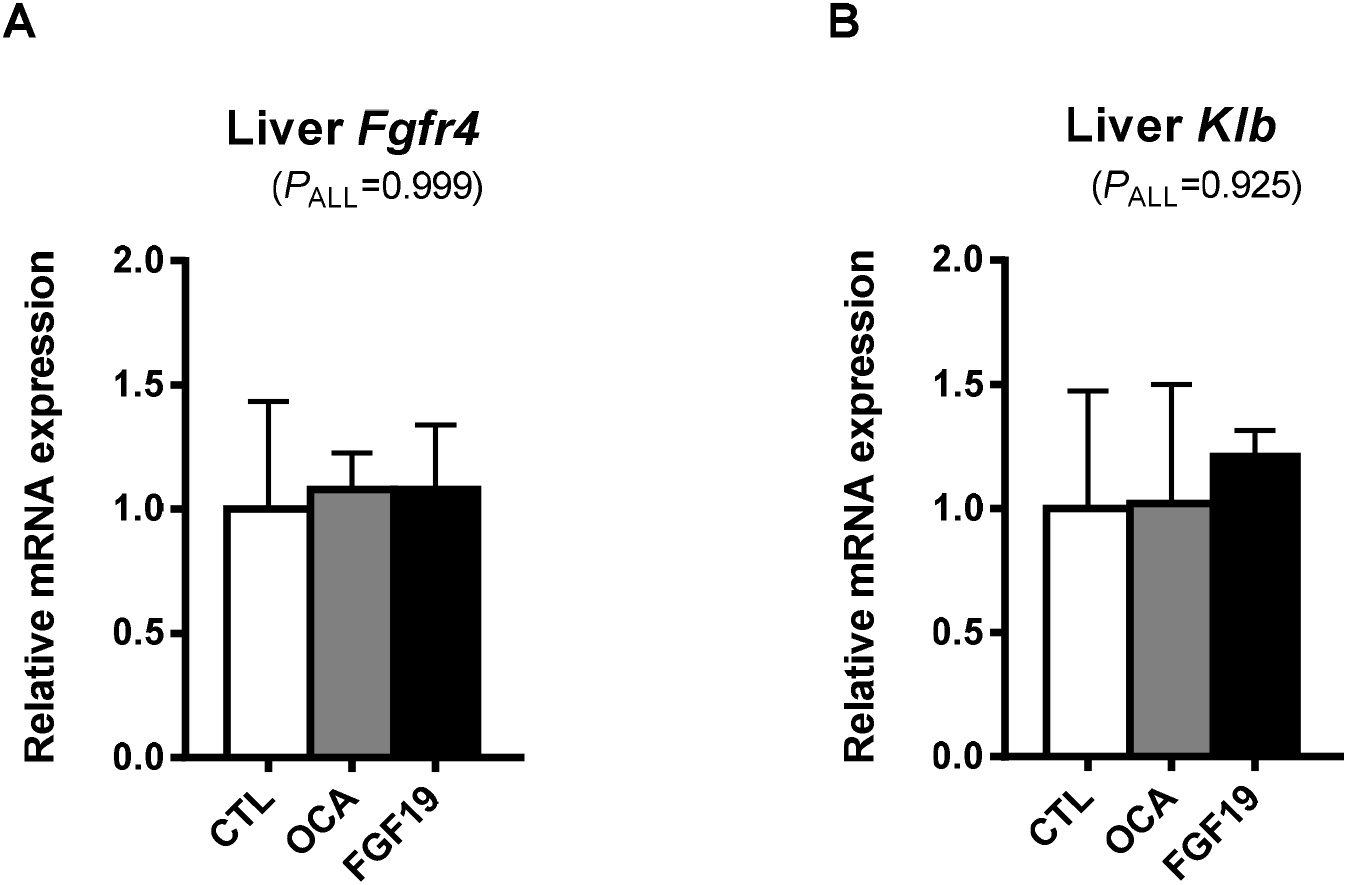
Activation of hepatic and ileal Fxr-Fgf15 pathways after pre-treatment with OCA, FGF19 or vehicle for seven days. Mice (n=5 mice per group) were pre-treated for 7 days by daily administration of OCA, FGF19 or vehicle, before sacrifice. Transcripts were analyzed in the liver (A, B). Values are expressed relative to the median expression in the control group. Data are displayed as median with interquartile range. CTL, vehicle-treated mice; OCA, obeticholic acid; FGF19, fibroblast growth factor 19; Klb, beta klotho.

**Figure S2.**
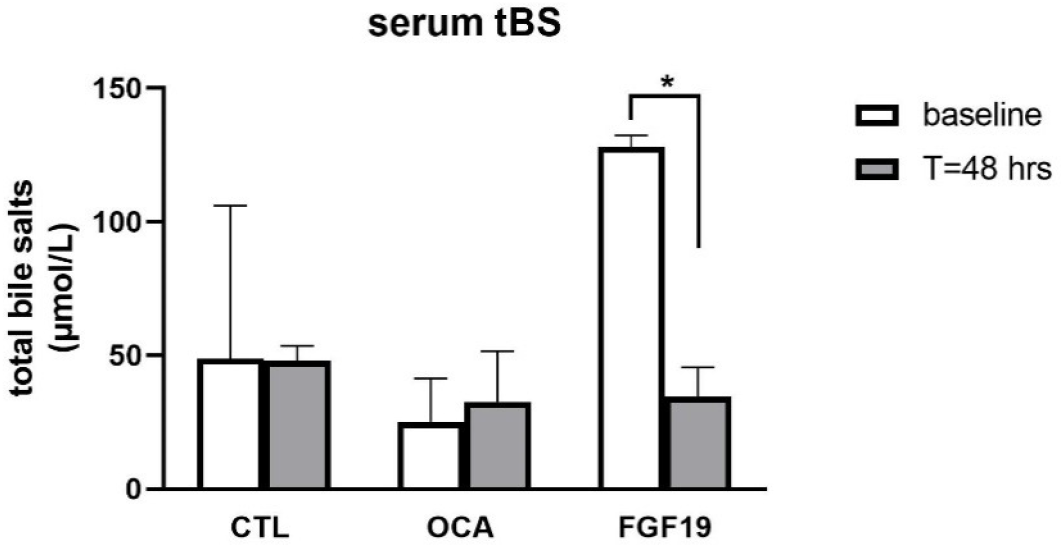
Serum total bile salt levels in groups of mice sacrificed without undergoing hepatectomy (baseline, n=5 per group), and at 48 hrs after 70% partial hepatectomy (n=8 or 16 per group). Data are displayed as median with interquartile range. *p<0.05. CTL, vehicle-treated mice; OCA, obeticholic acid; FGF19, fibroblast growth factor 19; tBS, total bile salt.

## Notes

**Conflict of interest:** None.

### Competing Interest Statement

The authors have declared no competing interest.

